# Differences in drug intake levels (high versus low takers) do not necessarily imply distinct drug user types: insights from a new cluster-based model

**DOI:** 10.1101/2024.07.15.603634

**Authors:** Diego Martinez Castaneda, Martin O Job

**Author notes:** Corresponding author: Martin O Job, Department of Biomedical Sciences, Cooper Medical School of Rowan University, 401 S Broadway, Camden, New Jersey, USA.

## Abstract

**Background:** Classifying psychostimulant users as high and low responders based on median split of drug intake levels has face-validity: these appear to be different types of drug users. However, because psychostimulant intake levels a) are defined by an inverted U-shaped dose response (IUDR) curve, and b) do not necessarily imply motivation for the drug, it is unclear that median split-designated high and low drug responders represent different drug user types.

**Aims:** To determine if median split-designated groups of high and low drug takers represent distinct groups when subjected to a new cluster-based model.

**Methods:** Male Sprague Dawley rats (n = 11) self-administered cocaine doses (0.00, 0.01, 0.03, 0.1, 0.3, 0.56 and 1.00 mg/kg/infusion) to reveal the IUDR curve per individual. We derived six variables defining the structure of the IUDR curve (amplitude, mean, width, and area under the curve: AUC) and the IUDR-derived economic demand curve (consumption at zero price or Q_0_ and the motivation for drug or α). We compared median split and clustering of all variables (cocaine dose, IUDR/demand curves) obtained.

**Results:** Median split of individual cocaine doses and IUDR curve-derived variables identified high versus low responders, but these groups were inconsistent with regards to group composition. Clustering of all cocaine doses revealed one cluster. Clustering of IUDR curve-derived variables revealed one cluster. Global clustering of all cocaine doses and all IUDR curve-derived variables revealed only one cluster.

**Conclusions:** High and low drug takers do not necessarily represent distinct drug user types.

## Introduction

There is face-validity to the idea that high drug takers and low drug takers represent different drug user types. For example, drug takers that consume higher levels of drug also tend to show higher drug seeking and higher likelihood of relapse after withdrawal (Edwards et al., 2007; Piazza et al., 2000; Sutton et al., 2000). According to DSM-V, a drug user is thought to have developed substance use disorders when the drug user consumes the substance despite negative consequences. For this reason, drug takers that consume higher levels of drug under punishment conditions are termed punishment-resistant/compulsive while those taking lower levels under the same conditions are termed punishment-sensitive/non-compulsive (Cadet et al., 2016, 2017, 2019; Campbell et al., 2018; Daiwile et al., 2024; Datta, Martini and Sun, 2018; Datta, Martini, Fan, et al., 2018; Domi et al., 2021; Duan et al., 2022; Durand et al., 2021; Giuliano et al., 2018, 2019, 2021; Hopf and Lesscher, 2014; Hu et al., 2019; Jayanthi, Ladenheim, et al., 2022; Jayanthi, Peesapati, et al., 2022; Krasnova et al., 2017; Marchant et al., 2018; Munoz et al., 2023; Pelloux et al., 2007, 2012, 2015; Subu et al., 2020; Sun and Yuill, 2020; Torres et al., 2017, 2018; Xue et al., 2012; Zhou et al., 2019). As such, under non-punished or under punishment conditions, high versus low drug responders are thought to represent distinct drug user types.

Apart from potentially identifying different drug user types based on whether they prefer high or low levels of drug intake, distinguishing phenotypes via other behavioral outcomes can also reveal that they have differential drug intake levels. For example, subjects that were already separated into high and low responders based on other drug-related behaviors were subsequently confirmed to express high and low responses, respectively, in their level of drug intake (Belin et al., 2011; Bush and Vaccarino, 2007; DeSousa et al., 2000; Gannon et al., 2021; Gulley, 2007; Kuhn et al., 2022; Mandt et al., 2008; Merritt and Bachtell, 2013; Nishida et al., 2016). Thus, high versus low drug intake levels may be an important behavioral outcome for understanding drug user typology and may lead to more individualized or group-centric options to address this health crisis.

To classify drug user types into high and low responders, the median split analysis of responses is typically employed. The median split categorizes responses above the median as high responses and responses below the median as low responses. However, median split analysis has been shown via several studies to be a problematic approach particularly with continuous data (Bissonnette et al., 1990; Cohen, 1983; DeCoster et al., 2009; Fitzsimons, 2008; Garcia et al., 2015a, 2015b; Gebhardt et al., 2014; Iacobucci et al., 2015; Irwin and McClelland, 2003; Iselin et al., 2013; Knüppel and Hermsen, 2010; Maccallum et al., 2002; MacCallum et al., 2002; Maxwell and Delaney, 1993; McClelland et al., 2015). Importantly, the results from median split (high and low responders) does not account for the possibility of moderate or intermediate responders as a distinct group on its own.

Apart from the problems with the median split itself, its use in drug user typology is also problematic for reasons that have to do with the dose of drug self-administered. Most studies that have categorized subjects into high versus low responders have done this at a single dose of the drug. If the relationship between drug intake levels and drug dose was linear then the subjects that are classified as high responders at a single dose would always be classified as high responders regardless of drug dose (Figure 1A). However, dose-response relationships for addictive substances like morphine, nicotine, and cocaine have been consistently demonstrated to follow a non-linear relationship, or an inverted U-shaped dose-response curve (IUDR curve), implying that a subject designated as a high taker may always be categorized as such for the entire IUDR curve if the curve of the high responders is shifted upwards and outwards (Figure 1B), but may not always be the case (Figure 1C-D). Because the dose of drug infused can affect drug intake levels in individuals in a non-linear manner, it is not clear if categorizing subjects into high versus low responders based on drug intake levels at a single dose and without accounting for the entire dose-response curve is sufficient to inform us if the drug users belong to different groups of users.

**Figure 1.**
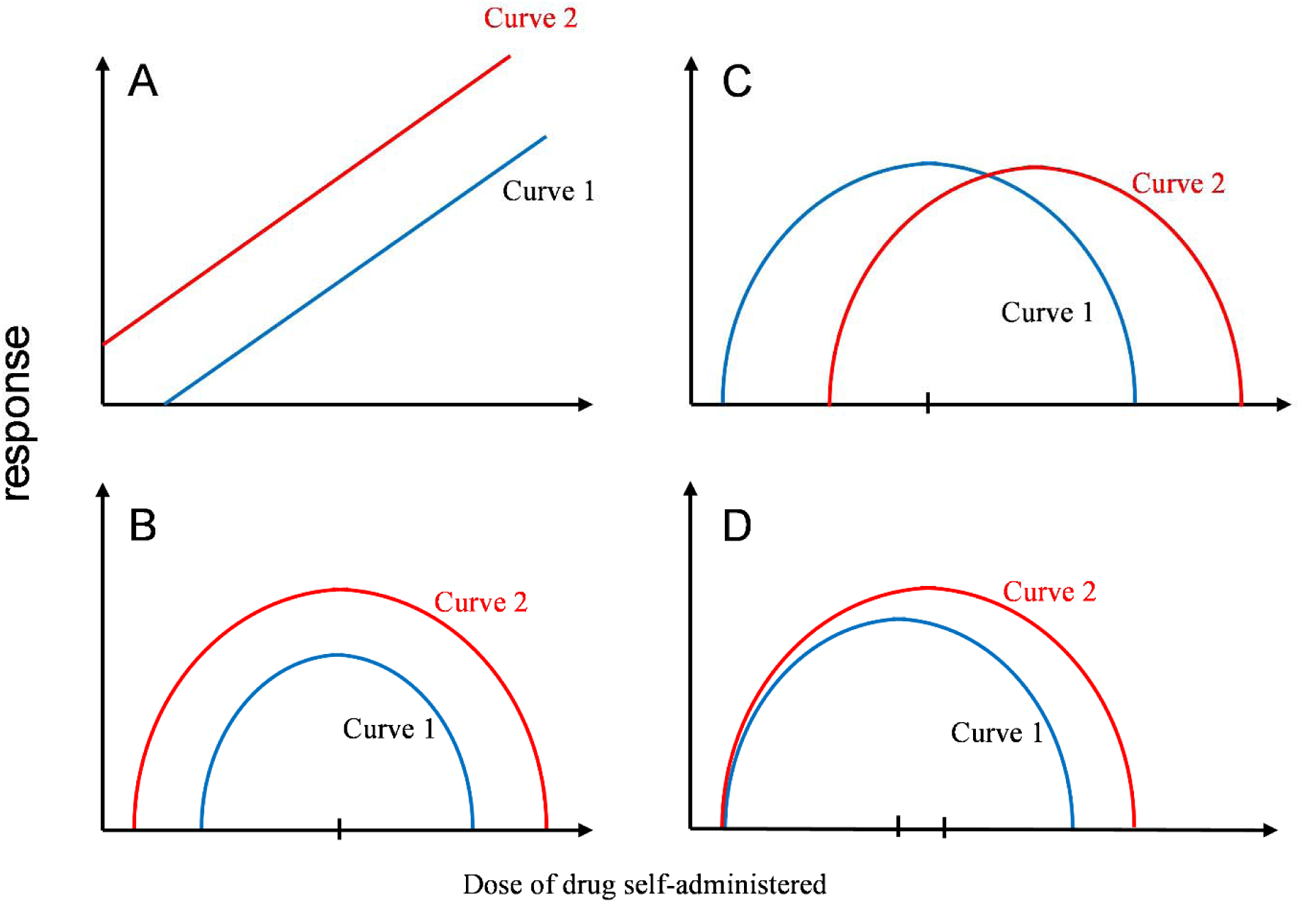
The problem of identifying high and low responders at more than one dose of an inverted U-shaped dose-response curve. In Fig A, the dose-response curve of two groups (curve 1 and curve 2) of drug users are both linear. The group designated curve 2 would always have a higher response compared to the group designated curve 1 at all doses of drug self-administered. Therefore if curve 1 and curve 2 represent low and high responders, respectively, these designations would be stable across all doses. In Fig B, the dose response curves of both groups are in line with inverted U-shaped dose-response curves, but as in Fig A, curve 2 would always have a higher response compared to the group designated curve 1 at all doses of drug self-administered. As in Fig A, if curve 1 and curve 2 represent low and high responders, respectively, these designations would be stable across all doses. However in Fig C, if curve 1 and curve 2 represent low and high responders, respectively, these designations would not be stable across all doses: curve 2 would be designated low responders at lower doses and high responders at higher doses, and vice versa for curve 1. In Fig D, curve 2 would be designated high responders and curve 1 would be designated as low responders at higher doses, but it would be difficult to distinguish these groups at lower doses. Thus, based on the inverted U-shaped dose-response curves of drugs, it is difficult to be confident that low and high responders identified at a single dose would represent stable groups.

Subjects that were separated into high and low cocaine intake groups based on median split at a single dose of cocaine (0.5 mg/kg/injection) under fixed ratio 1 schedule of reinforcement (FR1 schedule) subsequently displayed slightly different IUDR curves for cocaine self-administration (Edwards et al., 2007) - the cocaine IUDR curve for high takers was shifted vertical and rightward, relative to the IUDR curve of low takers. Furthermore, when subjects were separated into high and low sucrose responders based on median split of sucrose intake over a test period and allowed to acquire amphetamine self-administration behavior, a subsequent comparison revealed that the IUDR curves of high sucrose responders was shifted vertical and rightward, relative to the IUDR curve of low sucrose responders (DeSousa et al., 2000). However, when high and low sucrose takers designated via median split were allowed to acquire cocaine self-administration behavior, subsequent IUDR assessments of cocaine IUDR curves revealed no differences between high and low sucrose takers under FR1 schedule (Gosnell, 2000). Interestingly, under progressive ratio schedule, which is thought to assess motivation for the drug, high sucrose responders increased cocaine intake relative to low sucrose responders at one dose of cocaine (Gosnell, 2000). Thus, when we differentiate high versus low responders (for reinforcers) prior to obtaining the IUDR curve, these group differences may be reflected in the IUDR curve structure of the drug self-administered and in motivation for the drug, but this is not clear.

That said, differences in the IUDR curve structure are not sufficient to imply differences in drug user type. For example, curve 2 is shifted to the right relative to curve 1 and curve 1 is shifted to the left relative to curve 2 (Figure 1C). These ‘shifts’ do not necessarily imply different drug user types but may be the same drug user type under different responses to pharmacological manipulation. For example, administration of certain pharmacological agents can shift the dose-response curve of cocaine rightward (Barrett et al., 2004; Brebner et al., 1999; Collins et al., 2012; Emmett-Oglesby et al., 1993; Kuzmin et al., 1996; Mantsch et al., 2007; Peltier and Schenk, 1991; Self et al., 1998; Thomsen et al., 2008; Zanettini et al., 2018) or leftward (Barrett et al., 2004; Caine et al., 1999; Hiranita et al., 2009, 2014; Kuzmin et al., 1996, 1997; Pentkowski et al., 2012; Roberts et al., 2003; Self et al., 1998; Smith et al., 2004; Zanettini et al., 2018). Additionally, certain pharmacological manipulations can shift the dose-response curve of cocaine downwards (Barrett et al., 2004; Caine et al., 1999; Collins and Kantak, 2002; Galaj et al., 2020; Goeders and Guerin, 2000; Mantsch et al., 2007; Mu et al., 2021; Pei et al., 2015; Smith et al., 2004; Tella, 1995; Yang et al., 2022; Zanettini et al., 2018). *These shifts in the IUDR curves are more akin to responses to pharmacological interventions but do not imply a change in drug user type*. Thus, if the IUDR curves for high and low responders are considered as shown in Figure 1, these groups may merely represent different responses to the drug, but not necessarily distinct drug user types, but this is not clear.

High intake levels relative to low drug intake levels do not necessarily mean high relative to low motivation for the drug. For example, in studies that have employed behavioral economic analysis of demand curves, the consumption of a drug was not necessarily related to the demand elasticity or motivation for the drug (Bentzley et al., 2013, 2014; Roberts et al., 2013; Siciliano and Jones, 2017) in the same way that appetitive and motivational components of reinforcer intake are not the same (Ikemoto and Panksepp, 1996; Oleson et al., 2011a). This means that, in theory, 1) subjects expressing high versus low drug intake may have similar motivation for the drug, and 2) subjects in the low drug taker group may be as motivated for the drug as a subject in the high drug taker group. If we utilized motivation as the metric to distinguish distinct drug user types, high and low drug responders with similar motivation for drug would belong to a single drug user type. Consequently, a drug user type may include a mixture of subjects that would be identified as high and low responders. Indeed, differences in the motivation for psychostimulants assessed using the progressive ratio schedule were sometimes observed between high and low responders (Bush and Vaccarino, 2007; Mandt et al., 2008), but not always (Gosnell, 2000). It is not clear that high and low drug takers have different motivations for the drug.

Thus, even though high versus low drug intake imply differences in drug intake levels and may represent different drug user types, there are some limitations to this current approach including: 1. the limitations of the median split *per se*, 2. the non-linear structure of the dose-response curve (the IUDR curve), and 3. motivation for the drug *per se*. All of these need to be carefully addressed before we can be confident in drug user typology based on high versus low intake levels.

To address the first limitation, we employed normal mixtures clustering analysis of more than one variable at the same time to determine if multiple variables from the same individual would identify what population/ cluster to which this individual belonged, relative to other individuals. Normal mixtures clustering is not saddled with the assumption that there are only two groups (high versus low)-there could be one group or there could be eight groups, and if so, median split approach may be a type of experimenter-imposed grouping bias. To address the second limitation, we developed a new quantitative approach to analyze the structure of the IUDR curve of individuals to obtain relevant variables that define this curve (Figure 2A). To address the third limitation, we assessed motivation for drug using behavioral economic analysis of economic demand curves that were derived from IUDR curves of individuals (Figure 2B) – this enables us to analyze IUDR intake and motivation at the same time. Our model involves the clustering of all IUDR curve-derived structural variables and all economic demand parameters. Because our cluster-based approach is not entrapped by the aforementioned limitations, we rationalized that it would reveal a better model for drug user typology. Thus, we hypothesized that median split of drug intake levels would result in high versus low drug takers, but these groups would not be distinct groups when subjected to our cluster-based model.

**Figure 2.**
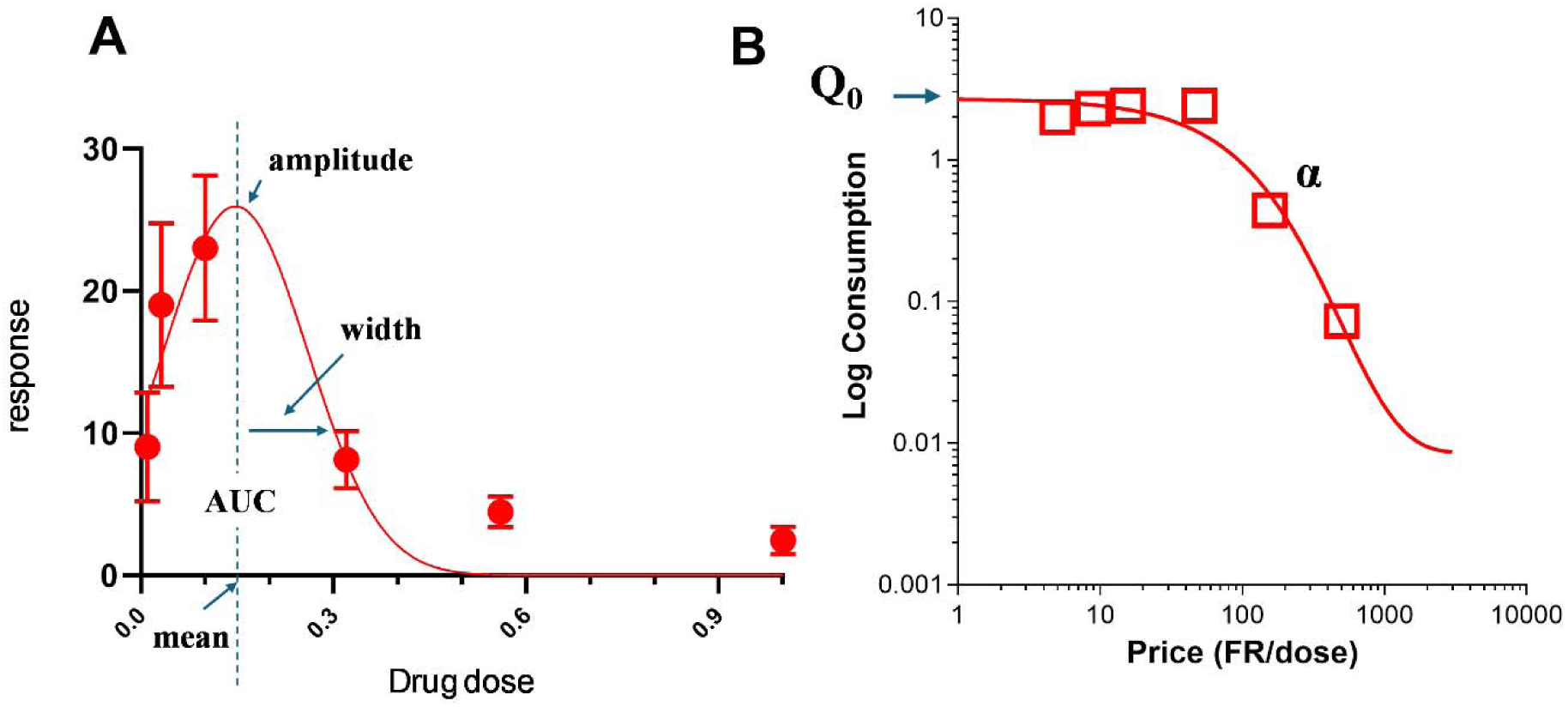
Derivation of variables from analysis of the structure of the inverted U-shaped dose-response (IUDR) curve (A) and behavioral economic analysis of the demand curves (B). Non-linear regression analysis of the IUDR curve using gaussian function (equation 1) to fit the curve yields the variables amplitude, mean, width and AUC (Fig A). From the IUDR curve, we can derive a demand curve with FR/dose on the x-axis and logarithm of responses/dose (amount of cocaine consumed) on the y-axis. From this demand curve, we can estimate the parameters Q_0_ - consumption at no cost and α – demand elasticity/motivation (Fig B).

To test our hypothesis, we used male Sprague Dawley rats. We allowed these subjects to self-administer multiple doses of cocaine to obtain the IUDR curve. We conducted median split analysis of responses at each dose to determine if respective individuals belonging to high versus low taker groups at a single dose would always be in the same designated groups for all doses. We compared the results from this to that obtained from clustering analysis of all the responses at all cocaine doses at the same time. Also, we derived several variables that define the IUDR curve structure and the demand curves (see Figure 2A-B) and, similar to what we did with cocaine doses as variables, we conducted median split analysis of each IUDR curve-derived variable to determine if respective high versus low responders at each variable would always be high versus low responders at all variables. We compared the results from this to that obtained from clustering analysis of all the IUDR curve-derived variables at the same time. We also conducted a global clustering of all cocaine dose variables and IUDR curve-derived variables/ economic demand variables. Our methods, results and discussion are below.

## Methods and Materials

### Subjects

Eleven male Sprague Dawley rats were obtained from Charles River Laboratories (Wilmington, MA). They were housed under temperature- and humidity-controlled conditions, and they were maintained on a 12-h/12-h light/dark cycle (lights on at 7:00 AM and off at 7:00 PM). After acclimation to the animal housing facility for at least one week, weights of rats were maintained at approximately 350 g via food restriction. Care of the animals was in accordance with the guidelines of the National Institutes of Health and the National Institute on Drug Abuse Intramural Research Program Animal Care and Use Program. The facility in which the research was carried out was accredited by AALAC International.

### Surgery

Chronic indwelling catheters (RJV-1, SAI Infusion Technology, Lake Villa, IL) were inserted under anesthesia (Isoflurane) using aseptic techniques into the external jugular vein of the rats. One end of the catheter was placed in the vein and the other end was a backmount/external port (313-000BM-10-5UP, P1 Technologies, Roanoke, VA) that exited from the back of the rat enabling access for flushing or infusions. This exit was closed with a screw cap (C313CAC, P1 Technologies, Roanoke, VA). Meloxicam was administered as an analgesic after surgery. To minimize the likelihood of infection and the formation of clots or fibroids, the catheter was flushed daily with 0.2 mL of a sterile solution containing heparin (30.0 IU/mL) and enrofloxacin (5 mg/kg). Rats were allowed to recover from surgery for a minimum of 1 week before cocaine self-administration protocols were initiated.

### Self-Administration

The operant-conditioning chambers for self-administration are as described in a previous paper (Job and Katz, 2019). All subjects were initially trained in 1-h daily sessions to respond for a 20-mg sucrose pellet under a fixed ratio 1-response (FR1) schedule. The pellet deliveries were followed by a twenty (20) second time out (TO) during which there was no reinforcer delivery and during which all lights in the operant chamber were off. After the subjects had acquired and maintained this behavior, the response requirement was changed to a FR5 schedule. Thereafter the subjects were allowed to self-administer cocaine (0.32 mg/kg/infusion) in 2-h daily sessions until they acquired and maintained this behavior. Afterwards, still on the FR5 schedule, the daily sessions were divided into five 20-min components, each separated by a 2-min TO. Then the subjects self-administered five different doses of cocaine per daily session (within-session design) with a single dose self-administered per component. To cover all the doses of cocaine the subjects had ten daily sessions on the within session design where they administered in five 20-min components, in order, cocaine dose = 0.032, 0.1, 0.32, 0.56, 1.00 mg/kg/infusion followed by eleven daily sessions where they administered, in order, cocaine dose = 0.0, 0.01, 0.032, 0.1, 0.32 mg/kg/infusion. The same concentration of cocaine (1.78 mg/mL) was placed in the syringes and the doses above were delivered by changes in delivered volume. For cocaine dose = 0, there was no solution delivered.

### Variable estimations and calculations

For each subject, we obtained the IUDR curves. We estimated, using non-linear regression analysis (see equation 1), behavioral variables that defined the IUDR structure (amplitude, mean, width and AUC, Figure 2A). We obtained a total of six (6) variables from the IUDR curve structure and the IUDR-derived economic demand curve (Figure 2A-B).

Where a = amplitude, x_0_ = mean, b = width

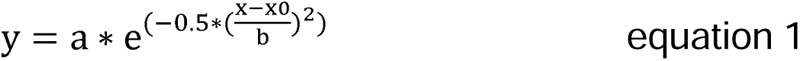

### Structural analysis of the IUDR curve

To obtain variables that define the IUDR curve structure, we employed nonlinear regression analysis of the IUDR curve using gaussian fit function (Figure 2A) as shown above.

### Behavioral economic analysis of the IUDR-derived economic demand curve

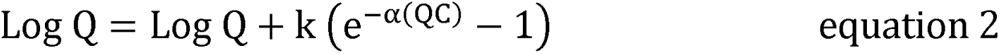

Where Q represents consumption of the reinforcer, C represents cost, Q_0_ represents consumption at no cost, α represents motivation and is a fitted parameter related to the decline in consumption with increased cost, and k is a scaling constant reflecting the consumption range.

From the IUDR curves, we obtained the demand curves. Demand curves were plotted with consumption (mg/kg) = number of reinforcers earned (infusions) × dose (mg/kg/infusion) on the y-axis and price = FR/dose (mg/kg/infusion) on the x-axis (Figure 2B). To obtain Q_0_ (consumption at zero price) and α (demand elasticity/ motivation which is related to the decline in consumption with unit increases in price), we employed behavioral economic analysis of the IUDR-derived demand curve using cocaine dose as a price (Figure 2B) (Bentzley et al., 2013; Oleson et al., 2011b; Oleson and Roberts, 2009; Yates et al., 2019). The demand curve was fitted using the exponential function (Hursh and Silberberg, 2008) (Figure 2B) as shown above.

Normal mixtures clustering analysis: We conducted normal mixtures clustering analysis of all subjects at 1) all responses for every dose of cocaine self-administered. We also conducted normal mixtures clustering analysis of 2) all variables derived from the IUDR curve and IUDR curve-derived economic demand curve. Finally, we conducted a global clustering analysis on all variables in 1) and 2), which included responses at multiple doses and IUDR curve structural variables and economic demand variables.

Median split analysis: We conducted median split of all subjects at all responses at each dose of cocaine self-administered. We also conducted median split analysis at the level of each variable derived from the IUDR curve and IUDR curve-derived economic demand curve.

Model comparison: We compared median split analysis and normal mixtures clustering analysis of the above referenced variables to determine if median split-designated high and low responders represented two distinct drug user types.

### Statistical analysis

We analyzed our data using the following software: GraphPad Prism v 10.2.3 (GraphPad Software, San Diego, CA), SigmaPlot 14.5 (Systat Software Inc., San Jose, CA) and JMP Pro v 17 (SAS Institute Inc., Cary, NC). We employed GraphPad templates for behavioral economic analysis/graphing from https://ibrinc.org/behavioral-economics-tools/. We employed Grubb’s test to identify and exclude outliers. For IUDR curve structure analysis for individuals, we employed gaussian fit of the IUDR curves to estimate amplitude, mean, width and AUC (see equation 1) – though not in the equation, we obtained AUC as an output from the analytical software together with the other variables (JMP Pro v 17). We employed multivariate analysis to determine the relationships between all the variables obtained. We employed unpaired t-tests to compare high versus low responder variable averages. For analysis of IUDR curve for median split-designated groups, we also employed 2-way repeated measures ANOVA to determine if there were any main effects of group and dose and if there were any interactions: group (high versus low) × cocaine dose (0, 0.01, 0.032, 0.1, 0.32, 0.56 and 1.0 mg/kg/infusion).

## Results

The Methods we employed are described above. Grubb’s test detected no outlier.

### Median-split designated group compositions were inconsistent across several cocaine dose self-administered variables

Table 1 shows all the subjects and their responses at each dose of cocaine self-administered. We conducted median split analysis of responses at each dose. The median value of responses per dose are shown at the bottom of the Table (Table 1). Based on values above and below the median, we designated groups as high (non-bolded) versus low (**bold**) responders, respectively, at each dose. The high and low taker groups were not consistent across all doses: some subjects that were designated as high takers at one dose were low takers at another dose (Table 1).

**Table 1:** Inconsistency in the group composition of high versus low responders across the dose response curve. Median values are shown in italics at the bottom row. Shaded values represent values below the median. The rat IDs (A-K) are shown in the first column. The numeric values are responses. The cocaine doses (including control – cocaine dose = 0) are in the 2^nd^ to 8^th^ column. Note that the group composition for high versus low responders varies with cocaine dose: depending on cocaine dose, every subject could be classified as both high and/or low taker. For example, rat A would have been designated a high responder at cocaine dose 0.1 and 0.32, but a low responder at cocaine dose 0.00, 0.01, 0.,032, 0.56 and 1.00.

| Cocaine |  |  |  |  |  |  |  |
| --- | --- | --- | --- | --- | --- | --- | --- |
| dose | 0.00 | 0.01 | 0.032 | 0.1 | 0.32 | 0.56 | 1 |
| ratID: A | **5.363636** | **7.272727** | **13.7619** | 24.09524 | 7.52381 | **4.1** | **2** |
| B | 5.545455 | **7.818182** | 24.66667 | 27.2381 | 8.238095 | 4.8 | 3.1 |
| C | **3.727273** | **6.181818** | 15.80952 | 27.66667 | 11 | 5.6 | 3 |
| D | 7.727273 | 9 | 30.04762 | **16** | **5.142857** | **3.2** | **1.5** |
| E | 9.636364 | 9.818182 | **15.33333** | 28.28571 | 11.2381 | 6.1 | 4.3 |
| F | **2.444444** | **2** | **5.5** | **14.61111** | **6** | 4.111111 | 2.777778 |
| G | **5** | **6.272727** | 17 | **16.04762** | **7** | **3.8** | **1.5** |
| H | **5** | 8.363636 | **15.71429** | **19** | **6.714286** | **3.5** | **1.7** |
| I | 8.545455 | 19.45455 | 24.7619 | **19.61905** | **6.142857** | **3.3** | **1.8** |
| J | 7.181818 | 8.363636 | 20.57143 | 22.7619 | 9.238095 | 4.4 | 2.4 |
| K | 6.363636 | 7.909091 | **12.42857** | 29.42857 | 9.047619 | 5.9 | 3.4 |
| median | 5.545 | 7.909 | 15.810 | 22.762 | 7.524 | 4.111 | 2.400 |

### Normal mixtures clustering of all responses at all cocaine self-administered doses for all subjects revealed only one cluster

We conducted normal mixtures clustering of all responses at every dose for every subject. We determined that there was only one cluster for all subjects at all cocaine doses self-administered (Figure 3A-B). Multivariate analysis (Figure 3C) revealed the following. Responses at cocaine dose = 0 were related to responses at cocaine dose = 0.01 (P = 0.0104) but unrelated to responses at cocaine dose = 0.032 (P = 0.0681), 0.1 (P = 0.4641), 0.32 (P = 0.6407), 0.56 (P = 0.8262) and 1.00 (P = 0.6472). Responses at cocaine dose = 0.01 were unrelated to responses at cocaine dose = 0.032 (P = 0.0566), 0.1 (P = 0.8252), 0.32 (P = 0.7431), 0.56 (P = 0.4704) and 1.00 (P = 0.6010).

**Figure 3:**
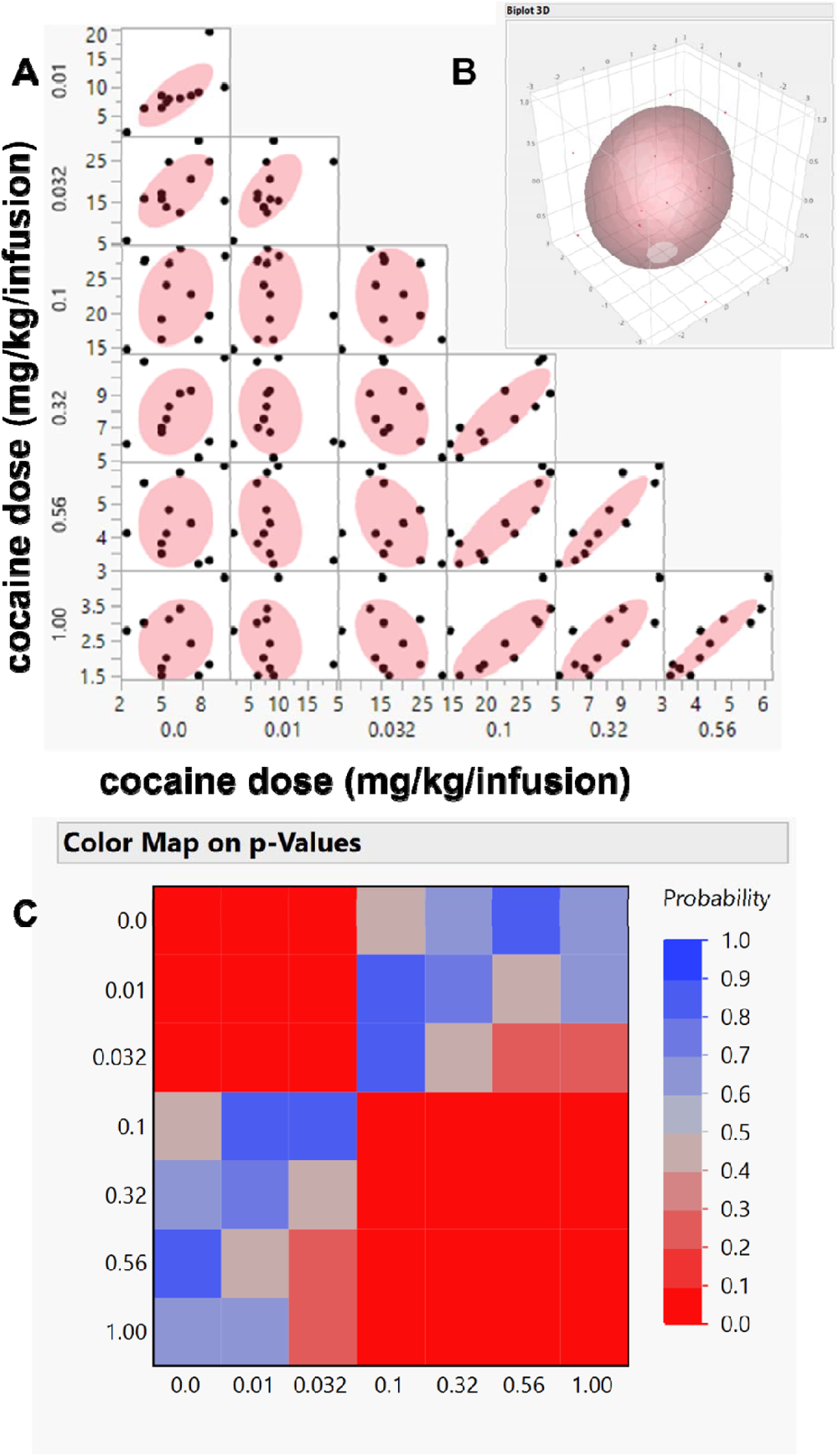
Normal mixtures clustering of all responses at all cocaine doses self-administered revealed that all subjects belonged to a single group. Fig A shows the results of the normal mixtures clustering. Fig B is a 3-D representation of Fig A. Fig C is the result of multivariate map of the analysis of all variables. For the multivariate mapping, deep red represents a P value < 0.05 and shows significant relationships between variables assessed. Deep blue represents a P value > 0.05 and shows that there are no significant relationships between variables assessed. Responses at higher cocaine self-administered doses were related.

Responses at cocaine dose = 0.032 were unrelated to responses at cocaine dose = 0.1 (P = 0.8437), 0.32 (P = 0.4844), 0.56 (P = 0.2434) and 1.00 (P = 0.2687). Responses at cocaine dose = 0.1 were related to responses at cocaine dose = 0.32 (P = 0.0012), 0.56 (P = 0.0009) and 1.00 (P = 0.0083). Responses at cocaine dose = 0.32 were related to responses at cocaine dose = 0.56 (P = 0.0002) and 1.00 (P = 0.0047). Responses at cocaine dose = 0.56 was related to responses at cocaine dose = 1.00 (P < 0.0001).

### Deriving the variables that define the IUDR curve structure and the IUDR curve-derived economic demand curve

For each subject, we employed gaussian function fit (equation 1) for the IUDR curves. From this model, we derived the amplitude, mean, width and AUC (see Figure 2A for variables, Figure 4 for individual subjects with R^2^ for goodness of fit). The (amplitude, mean, width and AUC) for subjects A, B, C, D, E, F, G, H, I, J and K were (30.61 ± 4.87, 0.1625 ± 0.0114, 0.09361 ± 0.00861, and 7.182 ± 0.918), (33.42 ± 9.62, 0.1522 ± 0.0250, 0.09826 ± 0.01856, and 8.232 ± 2.042), (39.17 ± 8.81, 0.1737 ± 0.0127, 0.09143 ± 0.01109, and 8.976 ± 1.463), (20.15 ± 8.31, 0.1051 ± 0.0667, 0.1142 ± 0.06316, and 5.766 ± 3.028), (35.02 ± 6.57, 0.1664 ± 0.0132, 0.102 ± 0.01207, and 8.955 ± 1.223), (23.45 ± 7.20, 0.1813 ± 0.0132, 0.08397 ± 0.01249, and 4.935 ± 1.047), (19.51 ± 6.04, 0.1544 ± 0.0276, 0.1141 ± 0.02718, and 5.582 ± 1.280), (22.6 ± 4.81, 0.1532 ± 0.0186, 0.1059 ± 0.01587, and 5.998 ± 1.016), (22.07 ± 5.40, 0.1063 ± 0.0417, 0.1281 ± 0.04178, and 7.085 ± 1.917), (27.53 ± 6.79, 0.1566 ± 0.0208, 0.1095 ± 0.01945, and 7.553 ± 1.401), and (39.57 ± 7.15, 0.1680 ± 0.0113, 0.08849 ± 0.00845, and 8.778 ± 1.244), respectively.

**Figure 4:**
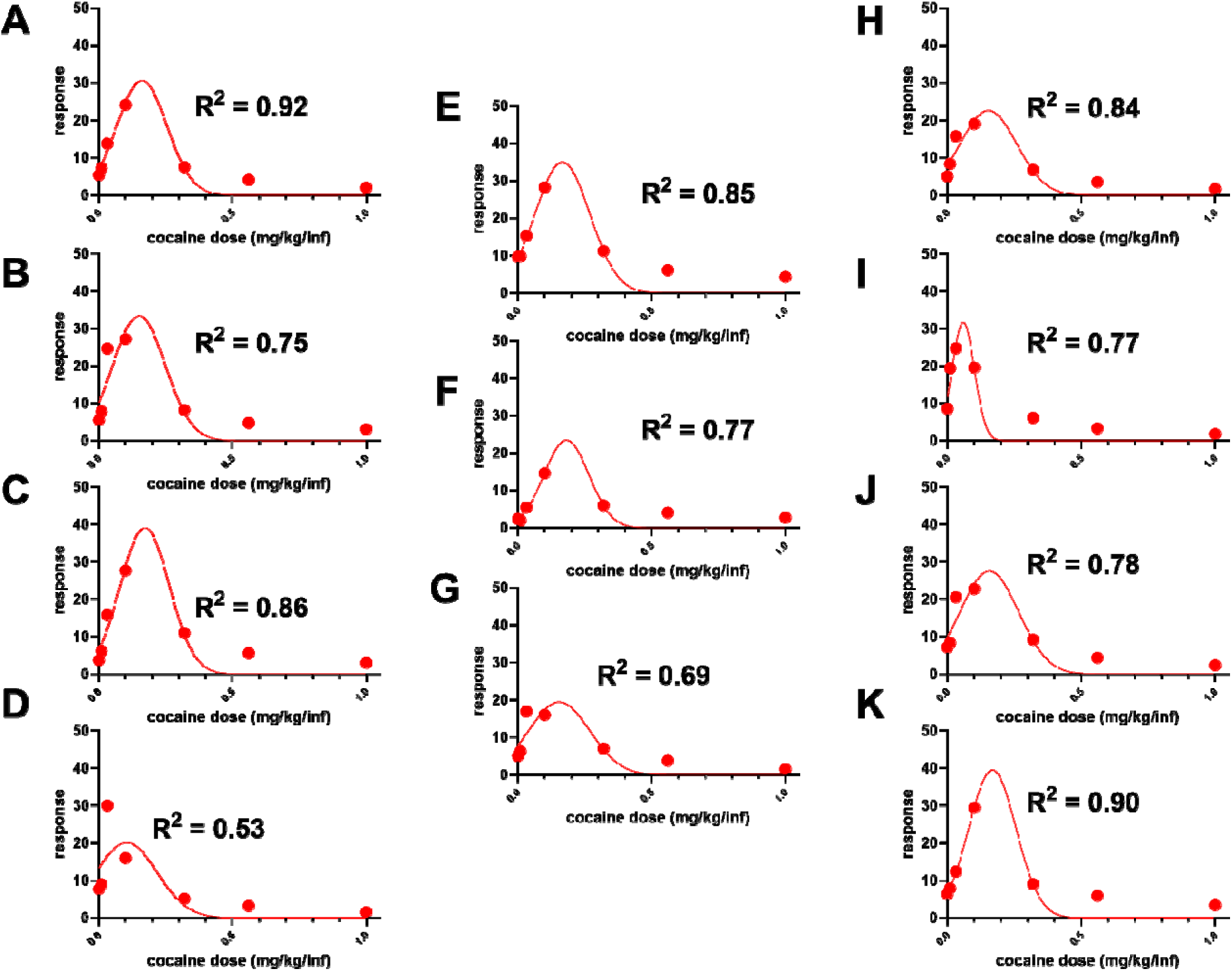
The inverted U-shaped dose-response curves for all individuals. A-J represent the inverted U-shaped dose response curves and the demand curves for individuals. R^2^ values (written into the graphs) for gaussian curve fit for IUDR curve for A-J were 0.92, 0.75, 0.86, 0.53, 0.85, 0.77, 0.69, 0.84, 0.77, 0.78, and 0.90, respectively. The amplitude, mean, width and AUC are shown in Table 2 and reported in the Results section.

**Table 2:** Inconsistency in the group composition of high versus low responders across several variables derived from the structural analysis of the IUDR curves and the behavioral economic analysis of the demand curves. Median values are shown in italics at the bottom row. Shaded values represent values below the median. Note that the high versus low group composition depends on the variable assessed: every subject could be classified as both high and/or low responder. For example, rat A would have been a high responder for amplitude, mean and α, but a low responder for width, AUC and Q_0_.

| Variables |  |  |  |  |  |  |
| --- | --- | --- | --- | --- | --- | --- |
| | amplitude | mean | width | AUC | $Q_0$ | $\alpha$ |
| ratID: |  |  |  |  |  |  |
| A | 30.61 | 0.1625 | **0.09361** | **7.128** | **2.7** | 0.00075 |
| B | 33.42 | **0.1522** | **0.09826** | 8.232 | 3.5 | **0.00059** |
| C | 39.17 | 0.1737 | **0.09143** | 8.976 | 4 | 0.00065 |
| D | **20.15** | **0.1051** | 0.1142 | **5.766** | **2** | 0.0007 |
| E | 35.02 | 0.1664 | 0.102 | 8.955 | 4.3 | **0.00055** |
| F | **23.45** | 0.1813 | **0.08397** | **4.935** | 3.1 | 0.0013 |
| G | **19.51** | **0.1544** | 0.1141 | **5.582** | **2.2** | 0.00084 |
| H | **22.6** | **0.1532** | 0.1059 | **5.998** | **2.2** | 0.00076 |
| I | **22.07** | **0.1063** | 0.1281 | **7.085** | **2.1** | **0.00051** |
| J | 27.53 | 0.1566 | 0.1095 | 7.553 | 3.1 | **0.00063** |
| K | 39.57 | 0.168 | **0.08849** | 8.778 | 3.9 | **0.00064** |
| median | 27.53 | 0.1566 | 0.102 | 7.182 | 3.1 | 0.00065 |

For each subject, we employed behavioral economic analysis using the exponential function (equation 2) for the IUDR curve-derived economic demand curve. From this model, we derived the Q_0_ and α (see Figure 2B for variables, Figure 5 for individual subjects with R^2^ for goodness of fit). The (Q_0_ and α) for subjects A, B, C, D, E, F, G, H, I, J and K were (2.70 ± 0.43 and 0.00075 ± 0.000093), (3.50 ± 0.38 and 0.00059 ± 0.00005), (4.00 ± 0.52 and 0.00065 ± 0.000062), (2.00 ± 0.29 and 0.00070 ± 0.000089), (4.30 ± 0.65 and 0.00055 ± 0.00006), (3.10 ± 0.38 and 0.0013 ± 0.00011), (2.20 ± 0.27 and 0.00084 ± 0.00008), (2.20 ± 0.26 and 0.00076 ± 0.000072), (2.10 ± 0.16 and 0.00051 ± 0.000038), (3.10 ± 0.30 and 0.00063 ± 0.000049), and (3.90 ± 0.71 and 0.00064 ± 0.000085), respectively.

**Figure 5:**
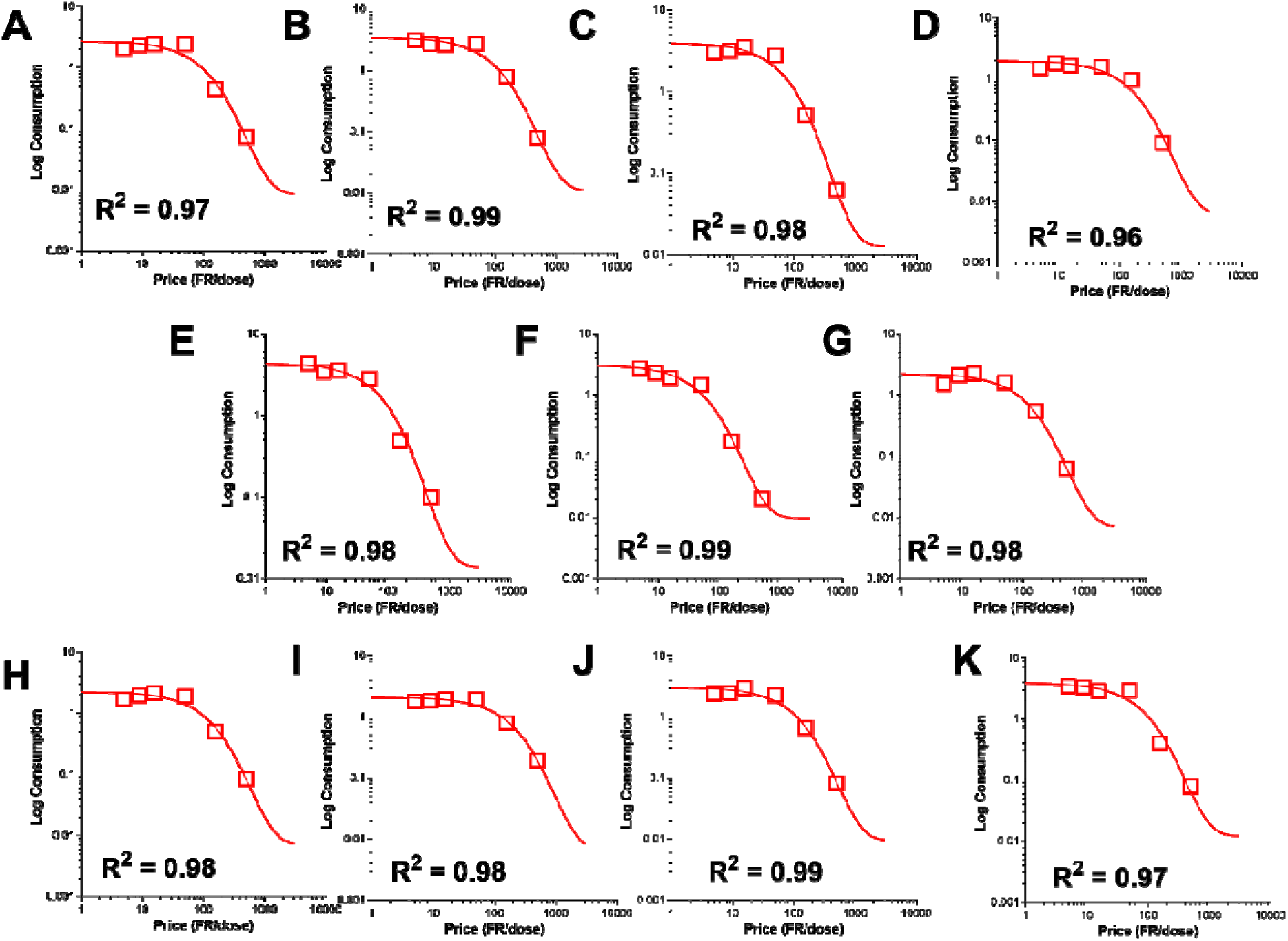
The economic demand curves derived from inverted U-shaped dose-response (IUDR) curves for all individuals. A-J represent the economic demand curves for individuals. R^2^ values (written into the graphs) for demand curves for A-J were 0.97, 0.99, 0.98, 0.96, 0.98, 0.99, 0.98, 0.98, 0.98, 0.99 and 0.97, respectively. The estimated Q_0_ and α are shown in Table 2 and reported in the Results section.

### Median-split designated group compositions were inconsistent across several IUDR curve-derived variables and demand curve-derived variables

Table 2 shows all the subjects and their variables with regards to the IUDR curve and demand curves. We conducted median split analysis of each variable and, based on the median value of each variable (bottom of Table 2), we designated groups as high (non-bolded) versus low (**bold**) responders at each variable. Per variable, the median split-designated high and low responder groups were not consistent across all doses. For example, high and low responders for the variable width were not consistently high and low responders for the variable AUC.

### Median split of individual variables and the effects of this procedure on other variables

In Table 2, we conducted median split for every variable and determined that this procedure yielded inconsistent groups. Next, we conducted median split at only one variable and examined the consequence of this procedure on the other variables.

Median split of consumption at zero price Q_0_ revealed high and low groups that were significantly different, as expected (P = 0.0003, Figure 6A). These groups were also different with respect to the variables-amplitude (P = 0.0164, Figure 6D), mean (P = 0.0378, Figure 6E), and width (P = 0.0438, Figure 6F). However, these groups did not differ with regards to α (P = 0.9172, Figure 6B). Furthermore, these groups were not different with regards to AUC (P = 0.0684, Figure 6G). The demand curve and the IUDR curve for high and low Q_0_ are shown in Figure 6C and Figure 6H, respectively. With factors group (high Q_0_, low Q_0_) and cocaine dose (0, 0.01, 0.032, 0.1, 0.32, 0.56 and 1.00), Two-way repeated measures ANOVA of the IUDR curve yielded a group × dose interaction (F 6, 54 = 3.371, P = 0.0068), a main effect of dose (F 2.285, 20.56 = 56.20, P < 0.0001), but no main effect of group (F 1, 9 = 0.1648, P = 0.6942). Sidak’s post hoc tests revealed that the IUDR curves of high versus low Q_0_ groups were significantly different for the highest cocaine doses (0.56 and 1.00 mg/kg/infusion).

**Figure 6:**
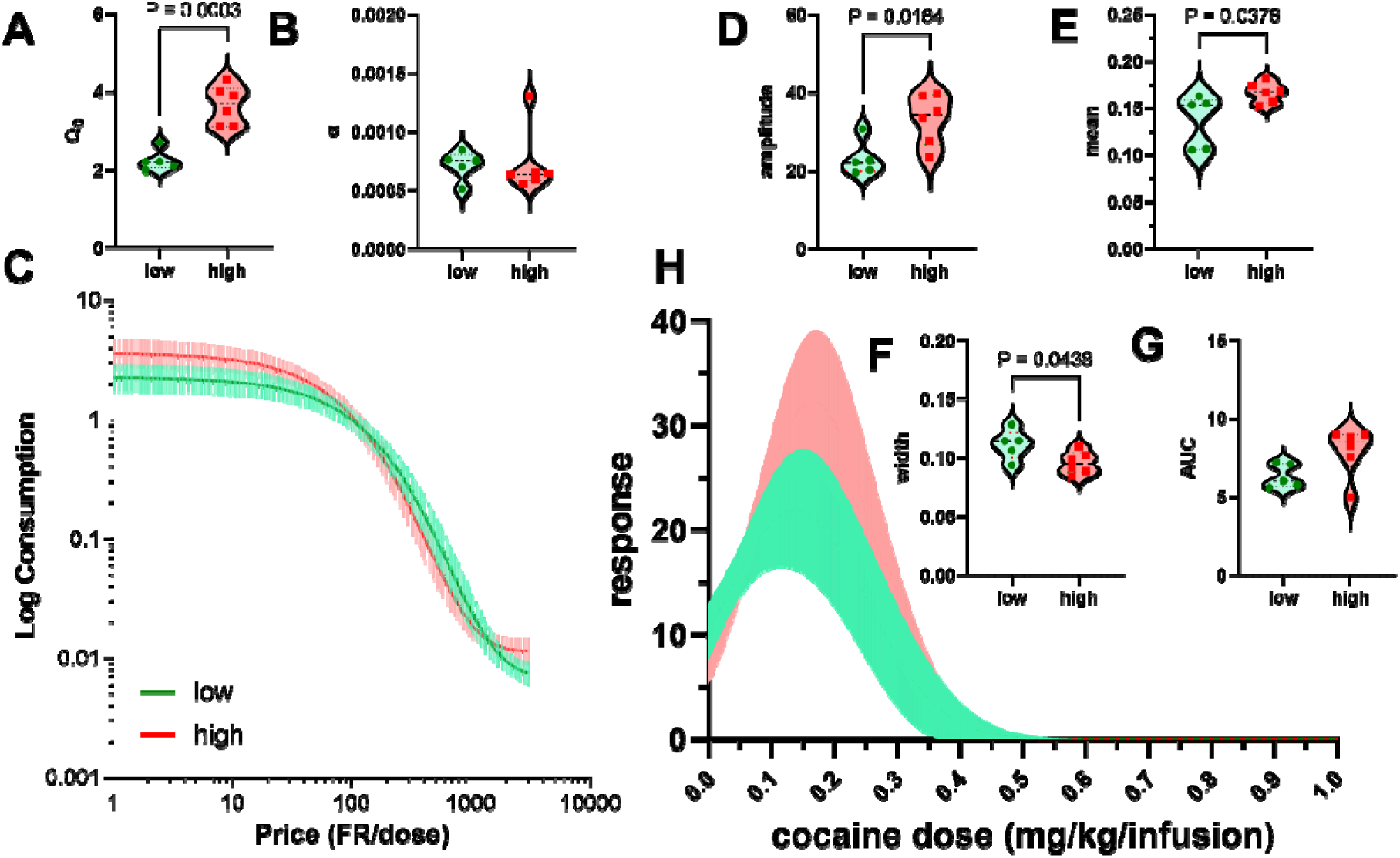
Median split of consumption at zero price Q_0_ revealed high and low groups but these groups were similar with regards to motivation for the drug (α). Fig A shows the result of median split of Q_0_ into high and low responders. Unpaired t-tests shows significant differences (P = 0.0003) between the groups above (high) and below the median (low). However, despite differences in Q_0_, these groups were not different with respect to α (P = 0.9172, Fig B). Fig C is a representation of the economic demand curves for the high versus low group. These groups were significantly different with regards to amplitude (P = 0.0164), mean (P = 0.0378) and width (P = 0.0438), but not AUC (P = 0.0684). In summary, low versus high groups showed differences in Q_0,_ but not α.

Median split of amplitude revealed high and low groups that were significantly different, as expected (P = 0.0003, Figure 7A). These groups were also different with respect to the AUC (P=0.0006, Figure 7A), but not mean (P = 0.1301, Figure 7A) and width (P = 0.1391, Figure 7A). For the IUDR-derived economic demand curve, these groups had significantly different Q_0_ (P = 0.0037, Figure 7B), but they were not significantly different with regards to α (P = 0.1609, Figure 7B). Out of a total of 6 variables, the median split-classification for high and low amplitudes were similar with regards to 3 variables (α, mean, width) and dissimilar with regards to 3 variables (amplitude, AUC, Q_0_). The implication was that the groups for high versus low amplitude were as similar as they were different (Figure 7C). The IUDR curve and the demand curve for high and low amplitude are shown in Figure 7A and Figure 7B, respectively. Two-way repeated measures ANOVA of the IUDR curve yielded a group × dose interaction (F 6, 54 = 3.782, P = 0.0032), a main effect of dose (F 1.671, 15.04 = 57.06, P < 0.0001), but no main effect of group (F 1, 9 = 2.981, P = 0.1183). Sidak’s post hoc tests showed that the IUDR curves that are revealed following median split to obtain high versus low amplitude groups were significantly different for the higher cocaine doses (0.1, 0.32, and 0.56 mg/kg/infusion).

**Figure 7:**
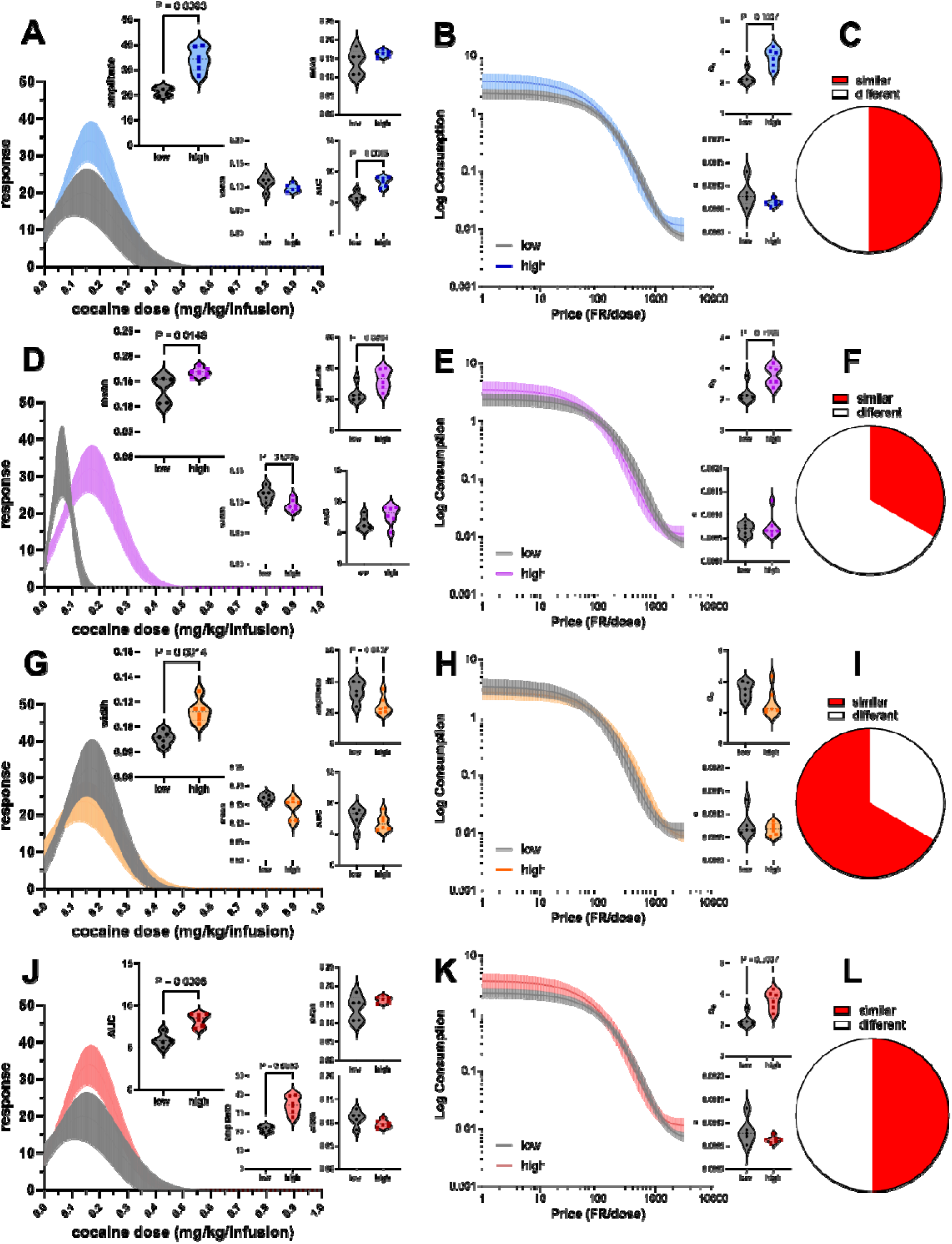
Median split of variables to identify high and low groups for amplitude, mean, width and AUC revealed distinct groups, but these groups were similar with regards to motivation for the drug (α). Fig A shows the result of median split of amplitude into high and low responders (P = 0.0003): these were also distinct with regards to AUC (P = 0.0006) and Q_0_ (P = 0.0037, Fig B insert), but not mean, width and α (P > 0.05). When we consider all 6 variables, high and low split by amplitude were as similar as they were different (Fig C). Fig D, G and J show the result of median split of mean, width and AUC, respectively, into high and low responders, but note that while these groups were significantly different with regards to some other variables (P <0.05), they were all similar with regards to α (P > 0.05). In summary, for all variables, median split revealed groups that while distinguished into high versus low responders, were all similar with regards to α (Fig B, E, H and K lower insert), while different with regards to some, but not all other variables assessed (Fig C, F, I and L).

Median split of mean revealed high and low groups that were significantly different, as expected (P = 0.0146, Figure 7D). These groups were also different with respect to the variables-amplitude (P = 0.0384, Figure 7D), width (P = 0.0206, Figure 7D), and Q_0_ (P = 0.0166, Figure 7E). However, these groups were not significantly different with regards to AUC (P = 0.1872, Figure 7D) or α (P = 0.6006, Figure 7E). Out of a total of 6 variables, the median split-classification for high and low mean were similar with regards to 2 variables (α, AUC) and dissimilar with regards to 4 variables (amplitude, mean, width, Q_0_). The implication was that the groups for high versus low mean were as similar for 33% of the variables assessed, but different for 66% of the variables assessed. They were not always different when all variables were considered (Figure 7F). The IUDR curve and the demand curve for high and low mean are shown in Figure 7D and Figure 7E, respectively. Two-way repeated measures ANOVA of the IUDR curve yielded a group × dose interaction (F 6, 54 = 6.269, P < 0.0001), a main effect of dose (F 2.934, 26.41 = 70.18, P < 0.0001), but no main effect of group (F 1, 9 = 0.09518, P = 0.7647). Sidak’s post hoc tests showed no difference between low versus high groups, at every dose.

Median split of width revealed high and low groups that were significantly different, as expected (P = 0.0014, Figure 7G). The IUDR curve and the demand curve for high and low width are shown in Figure 7G and Figure 7H, respectively. These groups were also different with respect to the amplitude (P=0.0457, Figure 7G), but not for any other variables - mean (P = 0.0672, Figure 7G), AUC (P = 0.3936, Figure 7G), Q_0_ (P = 0.1217, Figure 7H) and α (P = 0.3808, Figure 7H). Out of a total of 6 variables, the median split-classification for high and low groups based on width were similar with regards to 4 variables (α, mean, AUC, Q_0_) and significantly different with regards to 2 variables (width, amplitude). The implication was that the groups for high versus low width were mostly similar (Figure 7I). With factors group (high width, low width) and cocaine dose (0 - 1.00), Two-way repeated measures ANOVA of the IUDR curve yielded a group × dose interaction (F 6, 54 = 3.287, P = 0.0079), a main effect of dose (F 2.322, 20.90 = 56.30, P < 0.0001), but no main effect of group (F 1, 9 = 0.4733, P = 0.5088). Sidak’s post hoc tests showed no difference between low versus high groups, at every dose.

The median split of AUC revealed the same results as that obtained with amplitude (Figure 7J-L, compare with Figure 7A-C). The IUDR curve and the demand curve for high and low AUC are shown in Figure 7J and Figure 7K, respectively.

### The effect of median split of motivation for the drug (α) on other variables

One thing that was clear was that all groups, regardless of which variable was undergoing median split analysis, were similar with regards to α (Figure 6-7). These groups were also not always different when all other variables were accounted for. We wanted to understand what would happen to other variables if we conducted a median split of α (Figure 8).

**Figure 8:**
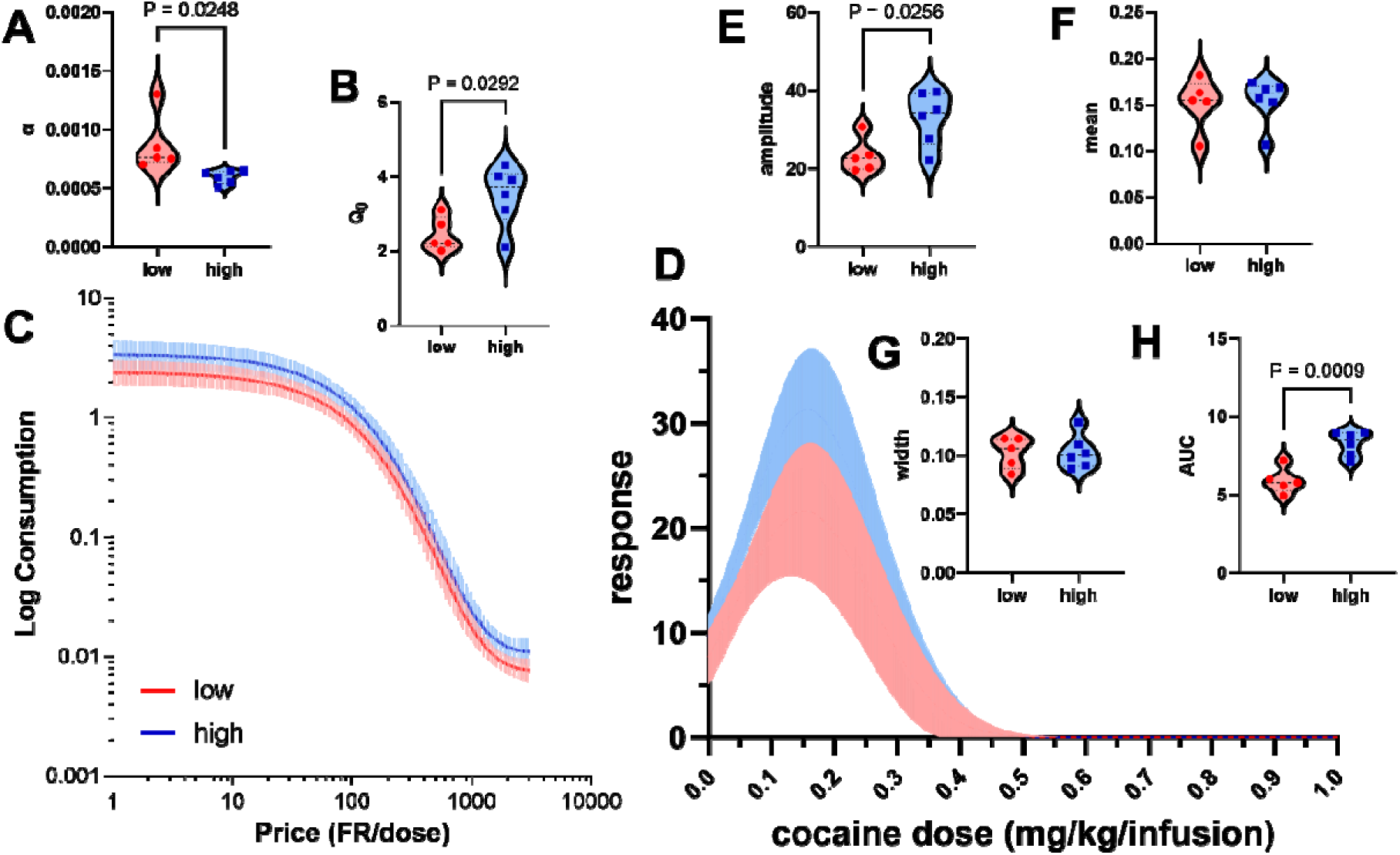
Median split of motivation for the drug (α) revealed groups that were different for some, but not all variables assessed. Fig A shows the result of median split of α into high and low responders. Fig B revealed that the resultant groups were also different with regards to Q_0_. Fig C shows the economic demand curve for the groups. Fig D shows the IUDR curves for the high and low α groups. Note that these groups were also significantly different with respect to amplitude (P < 0.05) and AUC (P < 0.05), but not mean (P = 0.8752) and width (P = 0.9442). In summary, even when we conducted median split of α, the groups corresponding to high versus low α were different for some, but not all variables assessed. Note that higher values of α represent lower motivation for the drug.

Median split of α revealed high and low groups that were significantly different, as expected (P = 0.0248, Figure 8A). The demand curve and the IUDR curve for high and low α are shown in Figure 8C and Figure 8D, respectively. These groups were also different with respect to the Q_0_ (P=0.0292, Figure 8B), amplitude (P = 0.0256, Figure 8E) and AUC (P = 0.0009, Figure 8H). These low and high α groups were not significantly different with regards to mean (P = 0.8752, Figure 8F) and width (P = 0.9442, Figure 8G). High and low α groups were similar with regards to 2/6 variables (mean, width) and different with regards to 4/6 variables (α, amplitude, AUC, Q_0_). The implication was that, even for median split-grouping for high versus low α, these groups were not always different. Note that higher values of α represent lower motivation for the drug. Two-way repeated measures ANOVA of the IUDR curve yielded no group × dose interaction (F 6, 54 = 1.178, P = 0.3319), a main effect of dose (F 1.724, 15.51 = 45.38, P < 0.0001), and a main effect of group (F 1, 9 = 12.74, P = 0.0060). Sidak’s post hoc tests showed no difference between low versus high groups, at every dose.

### Normal mixtures clustering of all IUDR curve-derived variables and demand curve variables for all subjects revealed only one cluster

We conducted normal mixtures clustering analysis of all variables at the same time. We determined that there was only one cluster for all subjects for all IUDR curve-derived variables and IUDR curve-derived economic demand parameters (Figure 9A-B). Multivariate analysis (Figure 9C) revealed the following. Q_0_ was unrelated to α (P = 0.6148) but was related to amplitude (P = 0.0001), mean (P = 0.0227), width (P = 0.0301) and AUC (P = 0.0037). α was unrelated to amplitude (P = 0.2631), mean (P = 0.1992), width (P = 0.1505), but was related to AUC (P = 0.0147). Amplitude was unrelated to mean (P = 0.0783) but was related to width (P = 0.0409) and AUC (P = 0.0001). Mean was related to width (P = 0.0012) but not AUC (P = 0.4536). Width was unrelated to AUC (P = 0.4888).

**Figure 9:**
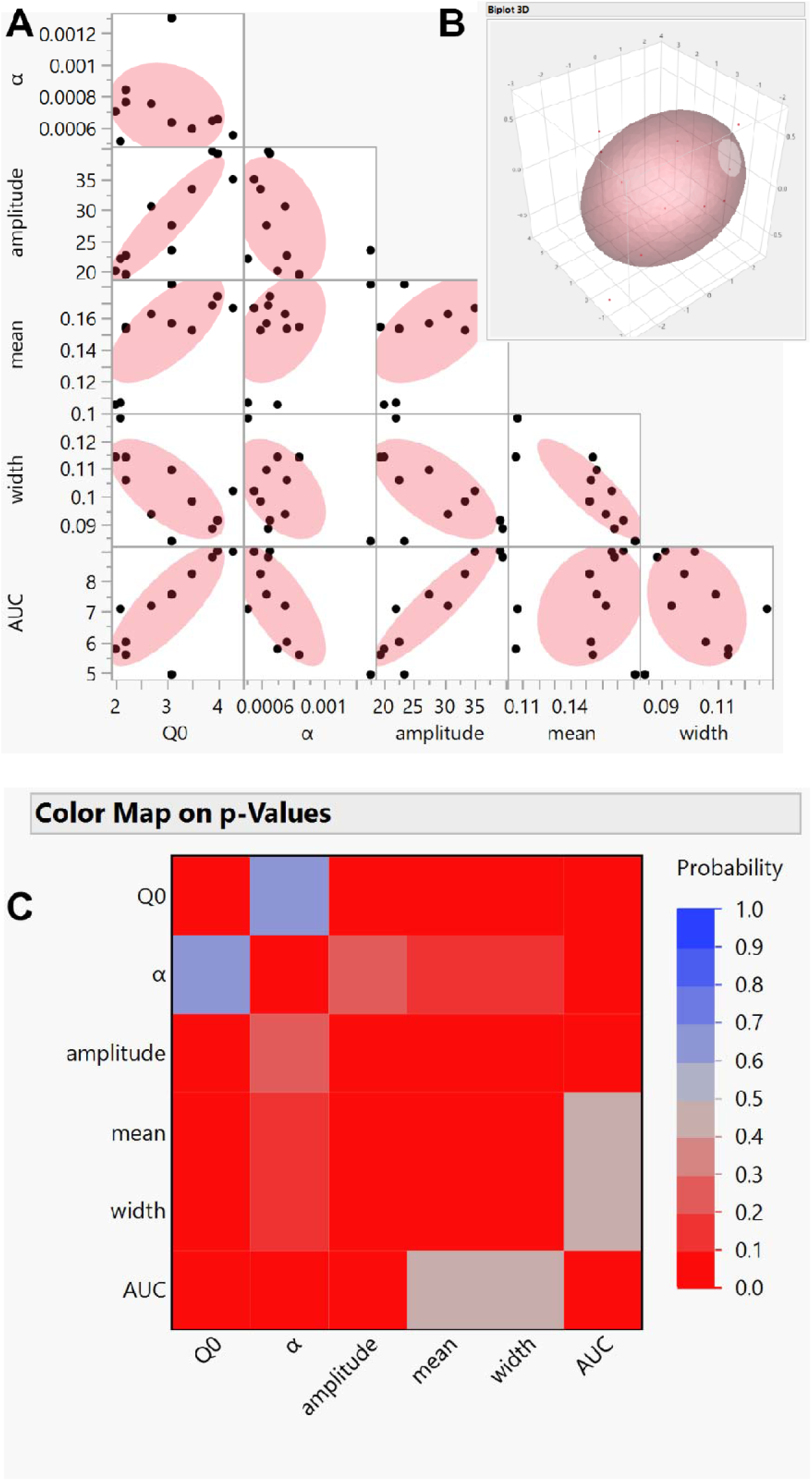
Normal mixtures clustering of all inverted U-shaped dose-response curve variables and demand curve variables revealed that all subjects belonged to a single group. Fig A shows the results of the normal mixtures clustering. Fig B is a 3-D representation of Fig A. Fig C is the result of multivariate map of the analysis of all variables. For the multivariate mapping, deep red represents a P value < 0.05 and shows significant relationships between variables assessed. Deep blue represents a P value > 0.05 and shows that there are no significant relationships between variables assessed. Normal mixtures clustering of the variables: Q_0_, α, amplitude, mean, width, AUC, revealed only one cluster. Q_0_ was related to amplitude (P = 0.0001), mean (P=0.0227), width (P = 0.0301) and AUC (P = 0.0037), but not α (P = 0.6148).

### Global clustering

We conducted normal mixtures clustering of all responses at all cocaine doses and all IUDR curve variables and economic demand variables. This revealed only one cluster. Multivariate maps of all variable relationships are shown in Figure 10. We have already reported the relationships between responses for cocaine doses (Figure 3) and between individual IUDR curve-derived variables. Here we report also the relationships between responses at cocaine doses and IUDR structure and demand economic curve variables. These were as follows. Responses at cocaine dose = 0 were related to α (P = 0. 0098) and width (P = 0.0463) and approaching a significant relationship for mean (P = 0.0696) but were unrelated to Q_0_ (P = 0.9888), amplitude (P = 0.9877), and AUC (P = 0.2803). Responses at cocaine dose = 0.01 were related to α (P = 0. 0172), mean (P = 0.0095), and width (P = 0.0058), but were unrelated to Q_0_ (P = 0.3838), amplitude (P = 0.6444), and AUC (P = 0.5295). For cocaine dose = 0.032, responses were related to α (P = 0. 0363), mean (P = 0.0008), and width (P = 0.0107), but were unrelated to Q_0_ (P = 0.2271), amplitude (P = 0.4157), and AUC (P = 0.8809). Responses at cocaine dose = 0.1 were related to Q_0_ (P = 0.0023), α (P = 0. 0438), amplitude (P < 0.0001), and AUC (P < 0.0001), but were unrelated to mean (P = 0.2544), and width (P = 0.2402). For cocaine dose = 0.32, responses were related to Q_0_ (P = 0.0004), amplitude (P = 0.0014), and AUC (P = 0.0006), but were unrelated to α (P = 0. 1815), mean (P = 0.0675), and width (P = 0.2689). For cocaine dose = 0.56, responses were related to Q_0_ (P < 0.0001), amplitude (P < 0.0001), and AUC (P = 0.0017), mean (P = 0.0318), and approaching significance for width (P = 0.0545), but were unrelated to α (P = 0. 4438). For cocaine dose = 1.00, responses were related to Q_0_ (P < 0.0001), amplitude (P = 0.0029), and AUC (P = 0.0114), but not mean (P = 0.0641), width (P = 0.0721), and α (P = 0. 6269).

**Figure 10:**
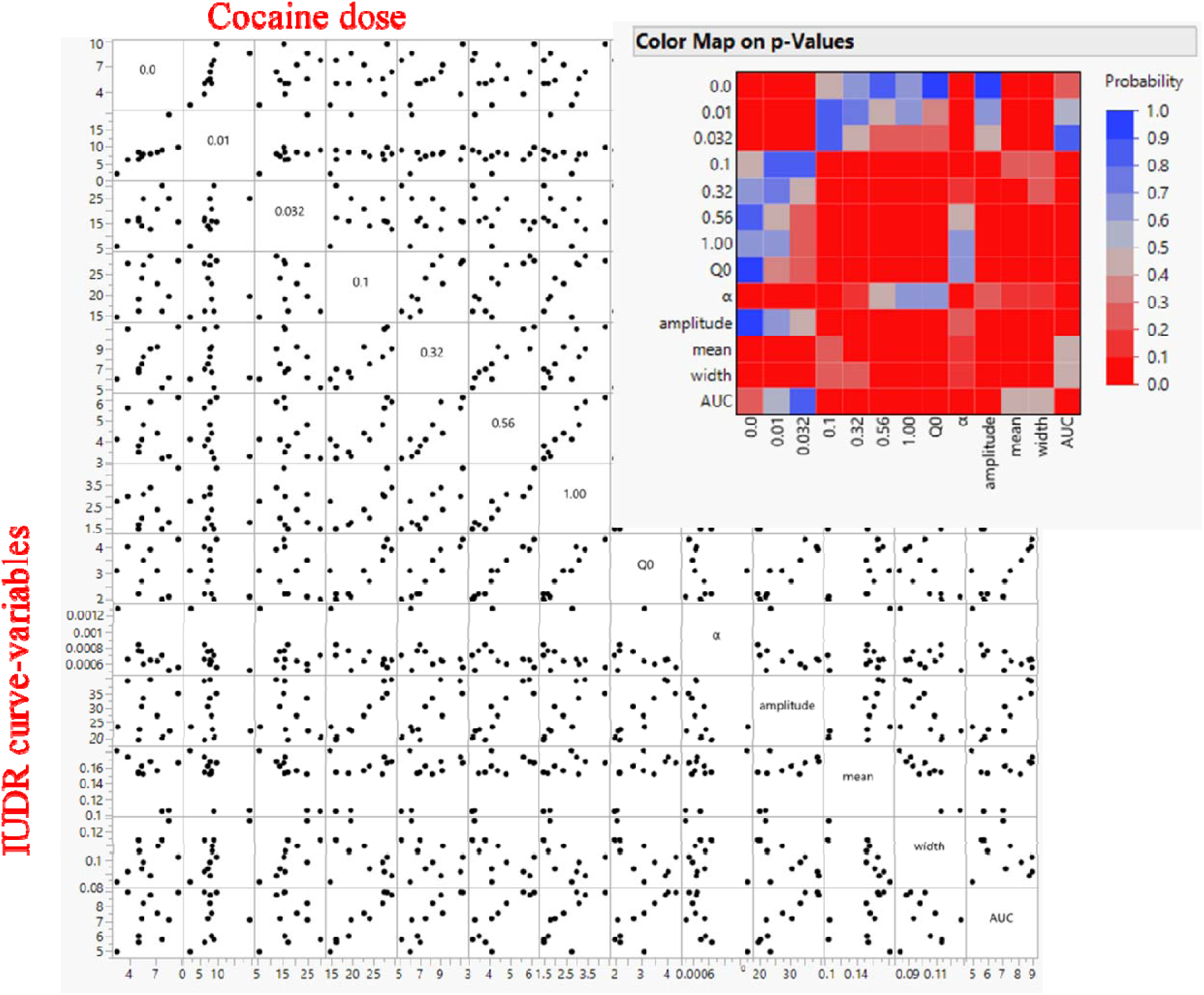
Normal mixtures clustering of all responses at all cocaine doses and all inverted U-shaped dose-response curve variables and demand curve variables revealed that all subjects belonged to a single group. The clustering analysis revealed only one group (data not shown). The multivariate graph and corresponding each map are shown for relationships between variables. For the multivariate mapping, deep red represents a P value < 0.05 and shows significant relationships between variables assessed. Deep blue represents a P value > 0.05 and shows that there are no significant relationships between variables assessed. Note that responses at higher cocaine doses are more related to Q_0_, amplitude and AUC whereas responses at lower cocaine doses are more related to α, mean and width.

### Summary of results

Median split analysis of responses at every dose of cocaine self-administered revealed high versus low responders, but these groups were inconsistent in composition across all doses – a high taker at a specific dose may not always be a high taker at another dose of drug self-administered (Table 1). Normal mixtures clustering analysis of responses at every dose of cocaine self-administered revealed only one cluster (Figure 3). Median split analysis of IUDR curve-derived variables for individuals (Figure 4-5) at the level of each variable also revealed high versus low responders at each variable, but these groups were inconsistent when we accounted for all variables (Table 2). Median split of individual variables revealed expected differences but the groups that emerged were not always distinct when other variables were considered (Figure 6-8). Normal mixtures clustering analysis of all IUDR curve-derived variables and IUDR curve-derived economic demand curve variables together revealed only one cluster (Figure 9). This was confirmed by global clustering of all variables that had to do with responses at all doses and all IUDR curve-derived variables relevant to IUDR curve structure and economic demand curves (Figure 10). There was only one group of subjects (Figure 3, 9, 10), not two. The data suggests also that high responses at higher doses did not necessarily imply higher motivation for the drug (responses were related to intake but not motivation at higher doses (Figure 10).

### Conceptualization for new criteria to distinguish drug user types

We determined that differential consumption, and differences in the IUDR curve variables were not necessarily associated with differences in α, suggesting that α may be the key variable required for distinguishing different drug user types. However, median split of α yielded groups that were not different at the level of all variables assessed, suggesting that α alone was not sufficient for drug user typology. Indeed clustering analysis revealed that there was only one drug user types detected in our data set. We propose that all six (6) variables, at least, are required to establish that two groups of drug responders are representative of different drug user types (Figure 11).

**Figure 11:**
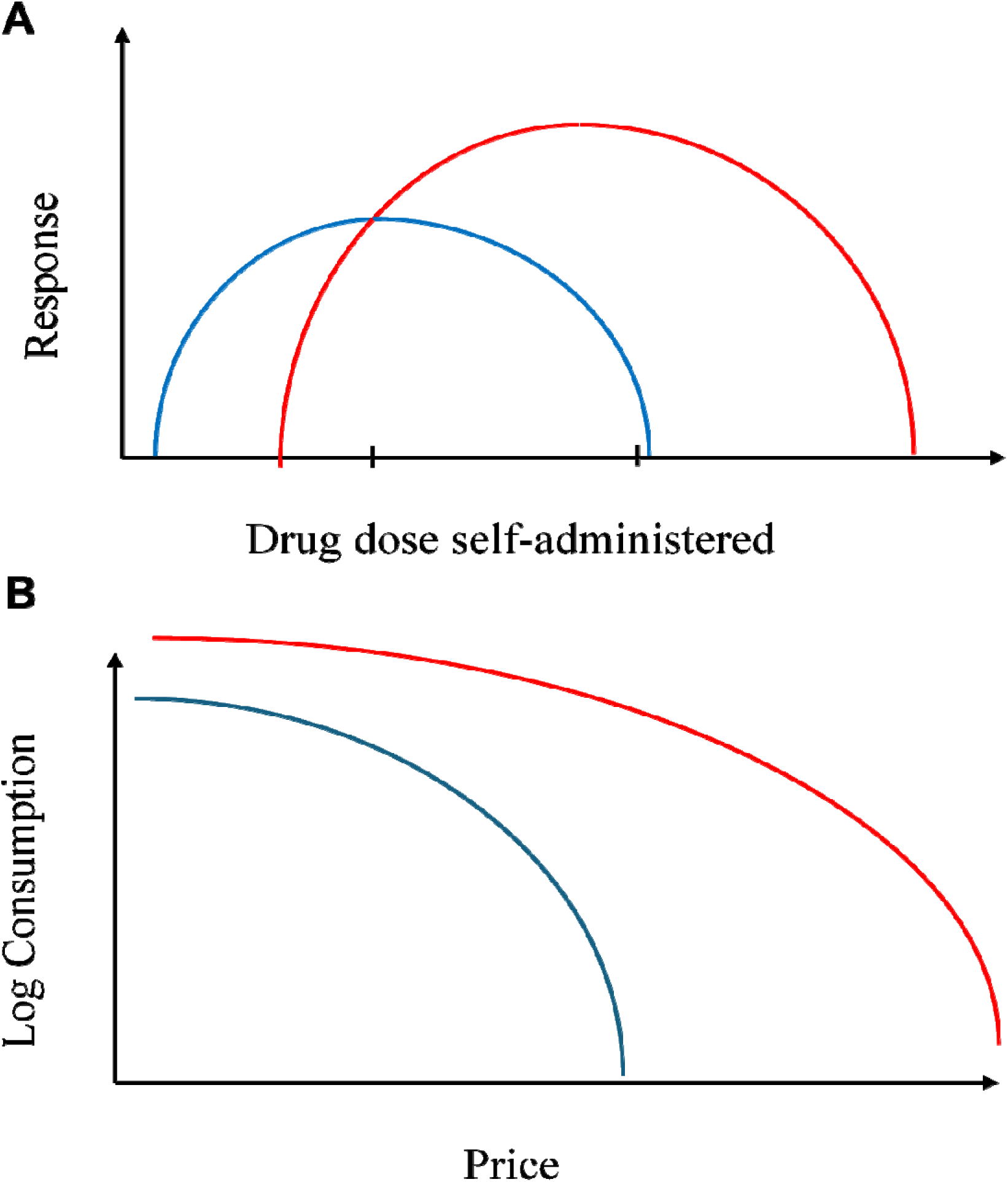
A standard model for determining distinct drug user types. Current models use median split analysis of one variable or variants of the median split. We show that while accounting for the IUDR curve and the economic demand curve, median split of any variable yields groups that are distinct on some, but not all other variables. By implication, these groups are not sufficiently different to give confidence that they belong to distinct drug user types. We propose that distinct drug user types will reflect significant differences in all the variables (amplitude, mean, width, AUC) of the IUDR curve and both variables (Q_0_, α) of the economic demand curve. This is like the groups identified in Fig7D-E but in addition, there is now a difference in α. This theoretical model will distinguish IUDR curves as shown in FigA and distinguish economic demand curves as shown in FigB.

## Discussion

Not every drug user develops substance use disorder, suggesting that there are different types of drug users. One approach employed in preclinical research to understand drug user typology is based on the idea that high versus low drug intake groups represent distinct drug user types. To conduct the separation of high versus low drug takers, the median split method is currently employed. However, the median split is usually conducted at a single selected dose of drug self-administered and may also be based on a single independent variable. This approach may be limited because the intake of self-administered drugs is defined by an IUDR curve, and high and low taker group compositions may vary with dose. Even when the IUDR curves of designated high versus low takers were obtained with differences noted, such as a vertical and rightward shift of the IUDR curves of high takers relative to low takers, it was/is unclear that these IUDR curve distinctions are sufficient to conclude that these groups are distinct drug user types. This is because in previous IUDR curves for high versus low takers (Edwards et al., 2007), there was an assumption, not a confirmation, that these groups also display different motivations for the drug. This assumption is an important question to answer because, as previously mentioned, motivation for the drug is not necessarily related to intake levels – we do not really know if the high takers are more motivated for the drug, we just know that they tend to consume more of the drug.

We developed a new approach to address the limitations of the current method. This new approach involved the clustering analysis of several variables including different cocaine doses, variables obtained from quantitative analysis of the IUDR structure and economic demand curve variables. These variables include intake and motivation. We hypothesized that median split of drug intake levels would result in high versus low drug takers, but these groups would not be stable when subjected to our cluster-based model. We predicted that the new approach would be more effective in identifying if high versus low takers represented distinct drug user types. We present data that confirms our hypothesis and highlights some major limitations of the current approach.

We determined that median split was not an effective tool in identifying drug user groups as it led to inconsistencies in group identification for different doses of self-administered cocaine and/or for different IUDR curve derived structural and economic variables. Even when we employed median split for individual variables, we determined that the high versus low responder groups that resulted were not sufficiently distinct. Clustering analysis, on the other hand, revealed only one group of responders based on cocaine dose and only one group of responders based on IUDR curve variables and economic demand variables. Global clustering – clustering of all cocaine dose and IUDR curve and economic demand variables – yielded only one cluster. Thus, our major finding is that our cluster-based model revealed that median split-designated high versus low drug takers do not necessarily represent distinct groups.

Based on our findings, we propose conservative criteria for drug user typology: for drug users to be considered different types they must be different with regards to all the variables of the IUDR curves and the economic demand curves. Under our criteria, our subjects belong to a single cluster. Our criteria expand on criteria already proposed in the field. For example, previous studies have suggested that vertical shifts (and rightward shifts) in the IUDR curve serve as a distinguishing factor for drug user typology as it relates to high versus low drug responders (Ahmed and Koob, 1998; Edwards et al., 2007; Piazza et al., 2000). We agree that amplitude (vertical shift) and mean (rightward shift) should be different for drug user type distinctions. By employing quantitative models to analyze the IUDR curves for the first time, we can add details to the current knowledge. For instance, we now also propose that width and the AUC must be different. Several of these variables are related to the economic parameter Q_0_. Another study suggests that drug user typology be based on α, or motivation or demand elasticity (Aston and Cassidy, 2019; Bentzley et al., 2013, 2014; Bentzley and Aston-Jones, 2017; James et al., 2019; Job and Katz, 2019; Kohtz et al., 2022; Mohammadkhani et al., 2019; Newman and Ferrario, 2020; Porter-Stransky et al., 2017). We agree. Our findings support all of these, but what we have now done is to bring all these together to develop a standard model for drug user typology that is based not on distinctions at the level of one variable (drug intake levels) but on distinctions at the level of several other variables too.

Our study corroborates a previous study that employed a similar animal model (male Sprague Dawley rats) of cocaine self-administration in which they employed median split to separate cocaine takers into high versus low takers and observed, just as we did, that the IUDR curve of cocaine for high takers was shifted vertical and rightward relative to the IUDR curve of low takers (Edwards et al., 2007), see also (Piazza et al., 2000). However, the authors in that study describe this vertical and rightward shift as indicative of an ‘addicted’ phenotype compared to the low intake group that did not express this behavior (Edwards et al., 2007). While we agree that the high intake group expresses this behavior, because we also observed it when we compared the IUDR curves of low and high responders for Q_0_, we do not agree that this represents an ‘addicted phenotype’. What a vertical shift and a rightward shift may imply is just that-a shift in some variables of the IUDR curve. But despite these shifts these groups are still more similar than they are different when we account also for other variables that define the same IUDR. Except we observe a significant shift in all variables, the current observations are not sufficient to imply that these groups are distinct. Importantly, our high versus low drug taker groups, though distinct with regards to total drug intake, were not distinct with regards to motivation and some of the other variables. Moreover, these described shifts in the IUDR curves between these groups are reminiscent of the effects of pharmacological manipulation on the IUDR curve, and not changes in a drug user from one type to another. But even if our observations are suggesting that these shifts in IUDR curve variables for high takers relative to low takers are revealing distinct drug user types, cluster-based analysis of several variables, including global clustering, puts an end to this speculation. Cluster-based analysis suggests that high and low drug takers are not necessarily distinct drug user types.

There are some limitations to this study especially because only male subjects were used, and it is plausible that different effects may be found in females (O’Connor et al., 2022). We will study sex differences in the future.

There are widespread implications of our study. Preclinical approaches that are based on differential drug intake as a behavioral marker to separate drug users into distinct drug use types or ‘phenotypes’, using current methods, may need to be interpreted carefully based on the limitations that we have mentioned above and based on our findings.

In summary, while there is face-validity in assuming that high versus low takers represent different drug user types, this is not necessarily so. When we employed a new model that included clustering of several variables associated with drug intake, we determined that there was only one cluster when the current model would have informed us that there were two. Our model does not focus on one variable, but rather analyses multiple variables from individuals and the interactions of these variables, to find groups of individuals that belong to the same population, or not. Our new objective approach may be important in advancing the field of drug user typology.

## Acknowledgement

The authors wish to acknowledge Dr. Jonathan L Katz in whose lab most of the experiments were carried out. MOJ conducted the behavioral experiments. Both authors contributed to data analysis and to the writing of the manuscript. MOJ designed and conducted the behavioral experiments and statistical analysis. This work was funded by the Department of Health and Human Services/National Institutes of Health/National Institute on Drug Abuse/Intramural Research Program, Baltimore, MD, USA [grant #-DA000547]. This work was also supported by the Francis Lax Fund for Faculty Development at Rowan University. This work was also supported by startup funds from Rowan University, Camden, New Jersey.

## Disclosures

The authors declare no conflict of interests with respect to research, authorship and publication of this article.

## Notes

### Competing Interest Statement

The authors have declared no competing interest.

